# NSUN2-mediated mRNA m^5^C Modification Regulates the Progression of Hepatocellular Carcinoma

**DOI:** 10.1101/2022.06.08.495406

**Authors:** Dan Song, Ke An, Wen-Long Zhai, Lu-Yao Feng, Ying-Jie Xu, Ran Sun, Yue-Qin Wang, Yun-Gui Yang, Quan-Cheng Kan, Xin Tian

## Abstract

RNA modification affects many biological processes and physiological diseases. The 5-methylcytosine (m^5^C) modification regulates the progression of multiple tumors. However, its characteristics and functions in hepatocellular carcinoma (HCC) remain largely unknown. Here, we found that HCC tissues had a higher m^5^C methylation level than the adjacent normal tissues. Transcriptome analysis revealed that a major fraction of the hypermethylated genes participated in the phosphokinase signaling pathways, such as the Ras and PI3K-Akt pathways. The m^5^C methyltransferase NSUN2 was highly expressed in HCC tissues consistently. Interestingly, the expression of many oncogenes was positively correlated with the expression of *NSUN2*, including *GRB2, RNF115, AATF, ADAM15, RTN3*, and *HDGF*. Real-time PCR assays further revealed that the expression of the mRNA of *GRB2, RNF115*, and *AATF* decreased significantly with the depletion of *NSUN2* in HCC cells. Furthermore, NSUN2 could regulate the cellular sensitivity of HCC cells to sorafenib via modulating the Ras signaling pathway. Moreover, knocking down *NSUN2* caused cell cycle arrest. Our study demonstrated a vital role of NSUN2 in the progression of HCC.

## Introduction

Liver cancer has the sixth-highest incidence and the third-highest mortality in the world and seriously threatens human health [1]. The main type of liver cancer is hepatocellular carcinoma (HCC), which is a primary malignant tumor originating from liver epithelial tissue or mesenchymal tissue [2]. Various kinase-related signaling pathways are aberrantly activated in HCC, such as the RAS/RAF/MAPK/Erk pathway (Ras pathway) and the PI3K/PTEN/Akt/mTOR pathway (PI3K-Akt pathway) [3, 4]. The Ras pathway regulates the proliferation, apoptosis, and differentiation of HCC cells [5]. This kinase pathway recruits the GRB2/SHC/SOS complex and promotes the phosphorylation of Ras and Raf when the membrane surface receptor of epidermal growth factor receptor (EGFR) receives the stimulation signal. Then, a high level of phosphorylated Erk (p-Erk), as an activation marker, translocates into the nucleus and combines with other transcription initiation factors to promote oncogene expression [6]. As the first-line molecular-targeted drug for HCC, sorafenib can specifically inhibit RAF phosphorylation in the Ras pathway and plays an important role in inhibiting HCC cell proliferation and angiogenesis [7, 8].

The 5-methylcytosine (m^5^C) modification occurs in many types of RNAs, including mRNAs and non-coding RNAs. NOP2/Sun RNA methyltransferase (NSUN2) mainly catalyzes the formation of m^5^C as a writer protein [9], induces the differentiation of epidermal and neural stem cells [10, 11], and directly affects gene expression in viruses by regulating the splicing of HIV-1 RNA [16]. Modified RNAs are recognized by Y-box binding protein 1 (YBX1) and Aly/REF export factor (ALYREF). YBX1 and ALYREF promote mRNA stability [12] and nuclear translocation [13] as the readers of m^5^C. No m^5^C eraser has been identified yet, although some proteins are involved in m^5^C oxidation. For example, AlkB homolog 1 (ALKBH1) and Ten-eleven translocation (TET) family proteins have been identified as dioxygenases that catalyze the conversion of m^5^C to hm^5^C, which regulates RNA degradation and mitochondrial activity [14, 15].

As a critical RNA m^5^C catalytic enzyme, the functions of NSUN2 have been described in multiple types of cancer. NSUN2 affects the mRNA stability of the heparin-binding growth factor (HDGF) by catalyzing m^5^C modification in its 3’-Untranslated region (3’UTR), which promotes the pathogenesis of bladder cancer [16]. Additionally, NSUN2 is overexpressed in breast cancer (BRCA) and hypopharyngeal squamous cell carcinoma (HPSCC) [17, 18]. Pan-cancer analysis showed that NSUN2 is positively correlated with DNA copy number and mRNA expression, which are associated with poor prognosis [18, 19]. NSUN2 can regulate the m^5^C modification of H19 lncRNA and promote the occurrence and development of HCC by recruiting G3BP1 [20]. The m^5^C profiles of circular RNA and mRNA were discovered in HCC [21, 22]. However, the biological significance of NSUN2 and the characteristics of the m^5^C modification in HCC have not been fully investigated.

In this study, we analyzed the characteristics of the mRNA m^5^C modification in HCC tissues compared to that of the adjacent tissues by single-nucleotide resolution. The mechanism by which NSUN2 regulates the expression of multiple target genes was determined at the bioinformatic and experimental levels. We examined the effect of NSUN2 on regulating HCC cell sensitivity to sorafenib by affecting the activity of the Ras pathway. Additionally, down-regulation of NSUN2 in HCC cells arrested the cell cycle. The mechanisms of m^5^C regulated by NSUN2 were involved in the progression of hepatocellular carcinoma.

## Results

### The mRNA m^5^C is frequently hypermethylated in HCC tissues

To reveal the m^5^C modification features in HCC, we collected 20 pairs of HCC tumor samples and analyzed the transcriptome data (RNA-seq) and RNA bisulfite sequencing data (RNA-BisSeq). The m^5^C sites were enriched in mRNA in the HCC tumor tissues and the adjacent tissues (Figure S1A). By analyzing the RNA-BisSeq data from HCC tissues and the adjacent tissues, the distribution characteristics of the m^5^C modifications in HCC mRNAs were found to be enriched downstream of the translation initiation site in the mRNA coding sequence (CDS) region (**Figure 1**A). The distribution pattern was consistent with other mammalian cells previously reported [13]. The proportion of the m^5^C modification in different regions of the mRNA was statistically analyzed. The proportion of methylation sites covered in the 3’UTR, CDS, and 5’-untranslated region (5’UTR) was similar in cancer tissues and adjacent cancer tissues, and the CDS region contained the highest number of m^5^C sites (Figure 1B). A sequence frequency logo displayed an embedding feature of m^5^C sites in CG-rich environments (Figure S1B). Our RNA-BisSeq data identified 2,482 m^5^C sites in mRNAs with differential methylation levels (see Supplementary Table S1). Additionally, we found that mRNA m^5^C modification in HCC tissues was significantly higher than that in the adjacent tissues for the overall methylation level (Figure 1C). We found 1,548 and 934 sites for hypermethylated and hypomethylated, respectively. The ratio of hypermethylated sites in tumor tissues was 62.36%, and the ratio of hypomethylated sites was 37.63% relative to that in the normal tissues (Figure 1D). A heatmap showed the differential methylation level of the m^5^C sites between the adjacent tissues and HCC tissues (Figure 1E). In summary, the level of mRNA m^5^C is frequently hypermethylated in HCC.

**Figure 1.**
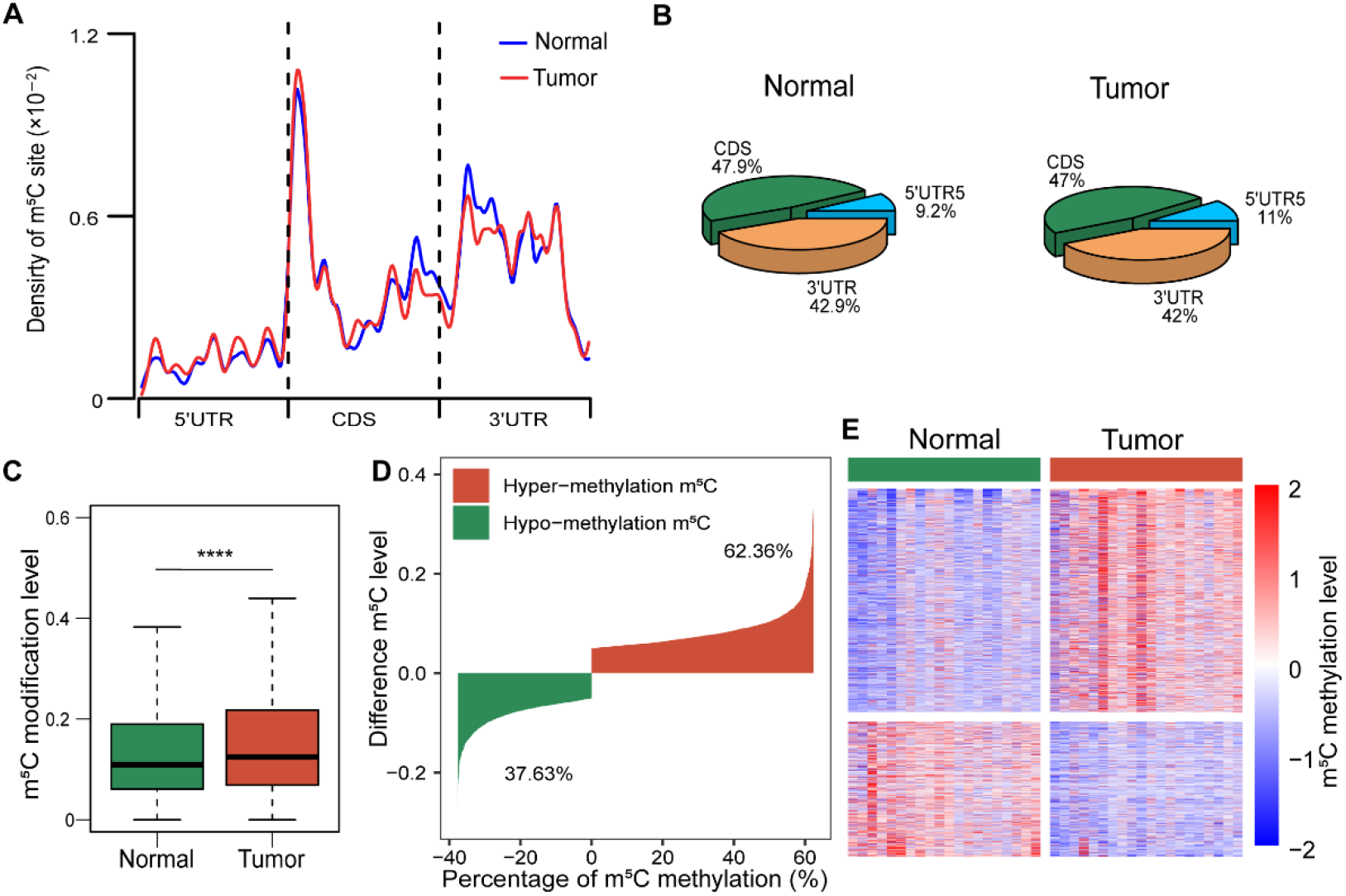
The mRNA m^5^C is frequently hypermethylated in HCC tissues. **A**. The distribution pattern of the m^5^C sites on mRNAs in HCC tissues (tumor) and the adjacent tissues (normal). **B**. Different proportion of m^5^C modification in regions of mRNA between HCC tissues and adjacent tissues. **C**. The overall m^5^C modification level was higher in HCC tissues than in the adjacent tissues, determined from the BisSeq data analysis. Statistical significance was calculated by wilcox.test, **** *P* = 5.116e−09. **D**. The difference in the m^5^C modification levels between HCC tissues and the adjacent tissues. **E**. A heatmap showing the differential m^5^C site methylation level between HCC tissues and the adjacent tissues. HCC, Hepatocellular Carcinoma.

### Multiple hypermethylated genes related to NSUN2 participate in carcinogenic pathways

To confirm these results and further understand the progress of HCC, we performed RNA-Seq and RNA-BisSeq in the same cohort of 20 HCC tissues and their corresponding normal tissues. We identified differentially expressed mRNAs with hypermethylated m^5^C sites in HCC tissues. We detected 255 hypermethylated sites, covering 124 genes with high mRNA expression (**Figure 2**A). Through the Kyoto Encyclopedia of Genes and Genomes (KEGG) analysis of highly m^5^C modified genes, several tumor-related pathways, including the PI3K-Akt, ErbB, and Ras signaling pathways, were found to be enriched in m^5^C modifications (Figure 2B). Moreover, these highly m^5^C-modified genes were found to be involved in the progression of cell migration, apoptosis, and cell cycle (Figure S1C). We investigated the genes (*GRB2, AATF, RNF115, ADAM15, RTN3*, and *HDGF*) that are highly expressed in HCC and modified by m^5^C (Figure S1D & S1E). To further determine whether the specific m^5^C modified genes are regulated by NSUN2 in HCC, first, we analyzed the correlation between the mRNA expression level of target genes and their m^5^C modification level (Figure 2C). Then, we determined the correlation between the *NSUN2* mRNA expression and the mRNA expression of the target genes (Figure 2D). The results showed that the mRNA expression of the target genes was related to their m^5^C modification level and also to the *NSUN2* mRNA expression level (Figure 2C & 2D). Additionally, the results of the TCGA analysis indicated that higher expression levels of these target genes (*GRB2, AATF, RNF115, ADAM15, RTN3*, and *HDGF*) were associated with a poor prognosis of HCC (Figure 2E & S1F). Thus, multiple hypermethylated genes associated with NSUN2 participate in oncogenic pathways.

**Figure 2.**
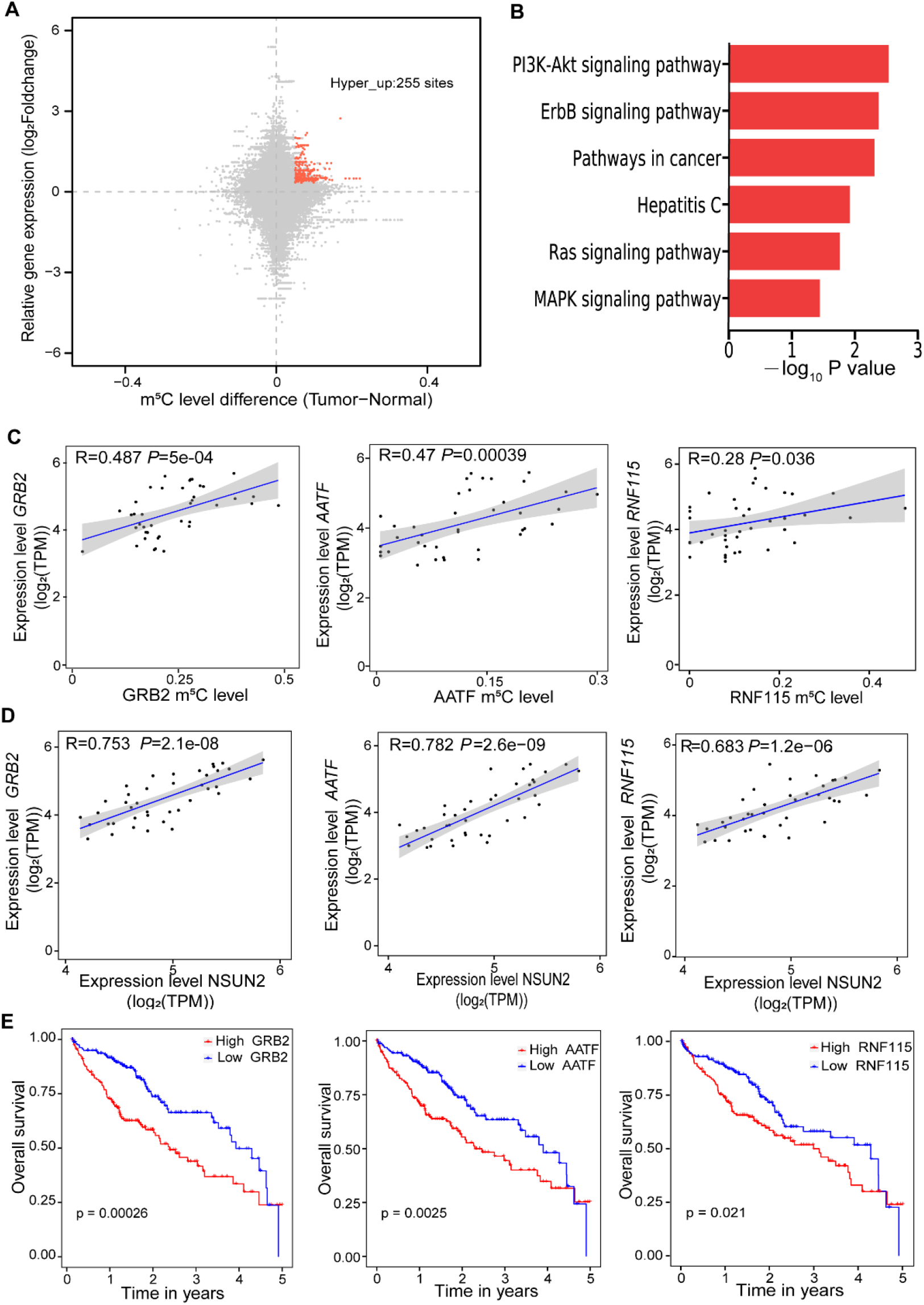
Multiple hypermethylated genes related to NSUN2 participate in the oncogenic pathways. **A**. The distribution of mRNAs with a significant change in the m^5^C methylation level and the gene expression level in HCC tissues and the adjacent tissues. **B**. The Kyoto Encyclopedia of Genes and Genomes (KEGG) analysis showed m^5^C hypermethylation and highly expressed genes in the HCC tissues enriched in oncogenic signaling pathways. **C**. A relation analysis showed that the expression levels of *GRB2, RNF115*, and *AATF* were positively correlated with their m^5^C modification levels; GRB2 (R = 0.487, *P* = 5e-04), RNF115 (R = 0.28, *P* = 0.036), AATF (R = 0.47, *P* = 0.00039). **D**. The expression levels of GRB2, RNF115, and AATF were positively correlated with the NSUN2 expression level; GRB2 (R = 0.753, *P* = 2.1e-08), NF115 (R = 0.683, *P* = 1.2e-06), AATF (R = 0.782, *P* = 2.6e-09). **E**. The overall survival analysis indicated the correlation of the mRNA expression of GRB2, RNF115, and AATF with poor prognosis in HCC patients; GRB2 (*P* = 0.0026), RNF115 (*P* = 0.021), AATF (*P* = 0.0025). Statistical significance was calculated by student’s t-test, mean ± SE.

### NSUN2 is highly expressed in HCC and regulates mRNA m^5^C modification

Transcriptome data analysis showed that m^5^C-related writer and reader proteins were highly expressed in HCC tissues (Figure S2A). Previous studies have shown that NSUN2 is involved in the regulation of m^5^C modification and affects tumor progression [20, 23]. Here we focused on the regulatory relationship between NSUN2 and m^5^C modified target genes in the progression of HCC. 20 pairs transcriptome data showed that NSUN2 mRNA was overexpressed in HCC tissues (**Figure 3**A). We also confirmed the protein expression of NSUN2 in some of the HCC tissue cohorts (n = 6) through western blotting (Figure 3B). Additionally, the results of the immunohistochemical analysis showed that NSUN2 had a higher expression in HCC tissues than that in the adjacent normal tissues (Figure 3C). Furthermore, the m^5^C modification level of total RNA and mRNA in the HCC cell lines (QGY-7703,SMMC-7721) were analyzed by ultrahigh-performance liquid chromatography-triple quadrupole mass spectrometry (UHPLC–MS/MS). We also used siRNAs to knock down NSUN2 and its family members (NSUN1/5). NSUN2 was found to be an important methyltransferase for mRNAs in HCC cells (Figure 3D & S2B). Real-time PCR revealed that the mRNA expression levels of *GRB2, RNF115*, and *AATF* were significantly decreased in the NSUN2 knockdown HCC cells (Figure 3E & S2C). The integrative genomics viewer (IGV) tracks displayed the read coverage of the GRB2 mRNA in the RNA-Seq and RNA-BisSeq and showed the upregulation of m^5^C and mRNA in HCC tissues relative to that in the adjacent tissues (Figure S2D). Therefore, we concluded that NSUN2 plays a critical role in regulating of the m^5^C modification of the target genes (*GRB2, RNF115*, and *AATF)* in HCC.

**Figure 3.**
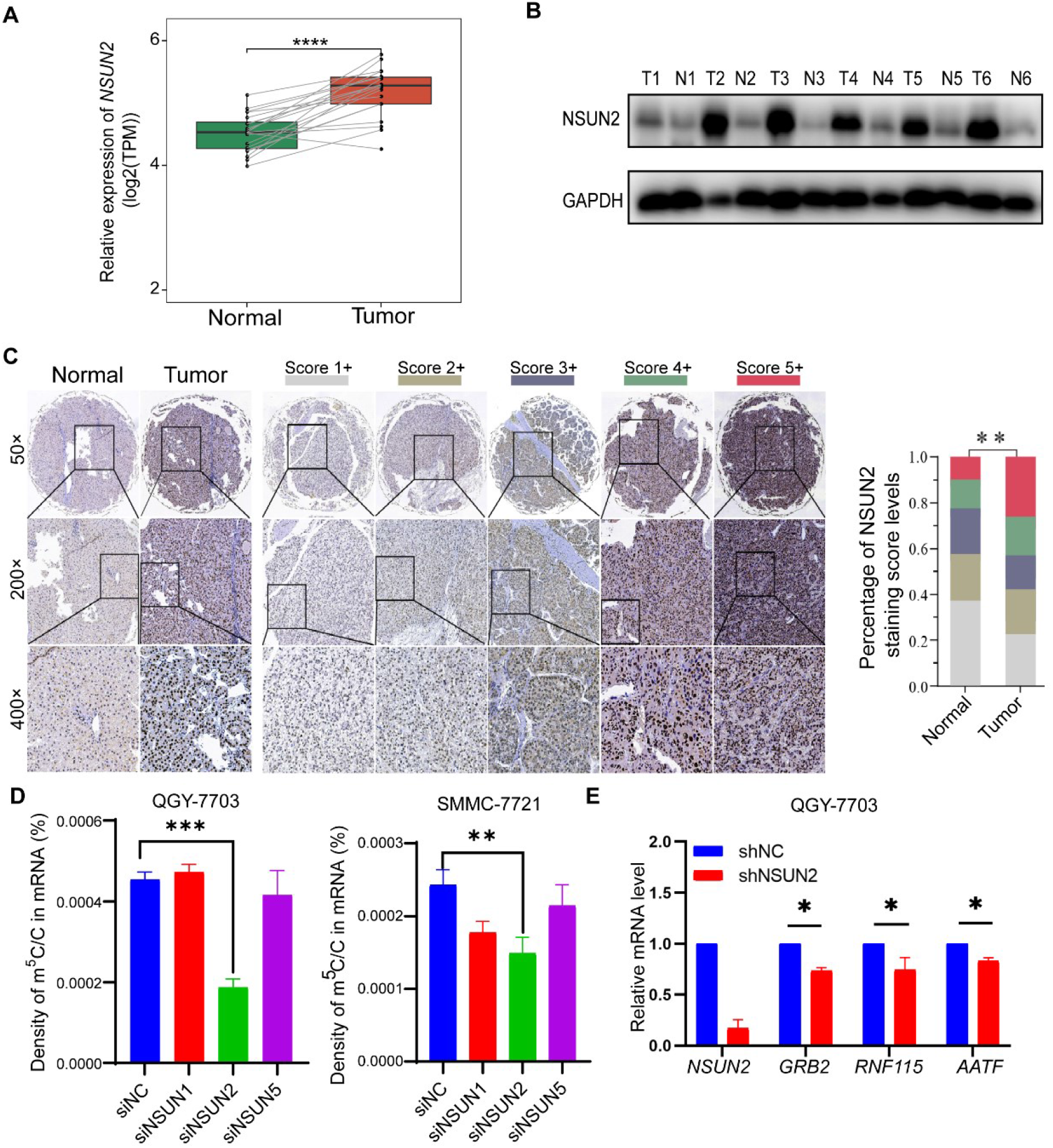
NSUN2 is highly expressed in HCC and regulates mRNA m^5^C modification. **A**. The expression of NSUN2 mRNA was higher in HCC tissues than in the adjacent tissues, determined by transcriptome analysis,\*\*\*\**P* < 0.0001. **B**. Western blot analysis showed that the expression of NSUN2 in HCC tissues was higher than that in the adjacent tissues. **C**. Immunohistochemical analysis showed high expression of NSUN2 in HCC tissues, \*\**P* < 0.01. **D**. In HCC cells, the UHPLC–MS/MS analysis showed that downregulation of NSUN2 significantly decreased the density of m^5^C/C in mRNAs (SMMC-7721, \*\**P* < 0.01. QGY-7703, *** *P* < 0.001.). **E**. The real-time PCR analysis showed that the mRNA expression of *GRB2, RNF115*, and *AATF* was significantly decreased when NSUN2 was silenced in GQY-7703 cells, \**P* < 0.05. Statistical significance was calculated by student’s t-test, mean ± SE.

### NSUN2 affects the sensitivity of HCC cells to sorafenib by regulating the activity of the Ras pathway

The activity of the Ras pathway is abnormally high in most HCC patients, which leads to a poor prognosis [24]. The phosphorylation of Raf is one of the crucial targets of sorafenib [25]. GRB2 is a critical upstream linker that promotes Raf phosphorylation that is regulated by NSUN2 in esophageal squamous cell carcinoma. Studies have shown that GRB2 is a key upstream regulator of Raf phosphorylation and is regulated by NSUN2 [26]. To determine the effect of the regulation of m^5^C by NSUN2 on the activity of the Ras pathway in HCC, the m^5^C modification levels of genes (such as *GRB2, MAPK3, PIK3R*) in Ras pathway were analyzed. These genes were hypermethylated in HCC tissues (**Figure 4**A). A heatmap analysis showed that the m^5^C modification level of these genes was increased in HCC tissues (Figure 4B). According to the results of the TCGA data analysis, HCC patients with higher expression of NSUN2 and GRB2 had the worst prognosis (Figure 4C). Since NSUN2 is important in regulating m^5^C modification and the activity of the Ras pathway, we further used the Ras activation assay kit to analyze the effect of NSUN2 on the level of active Ras.

**Figure 4.**
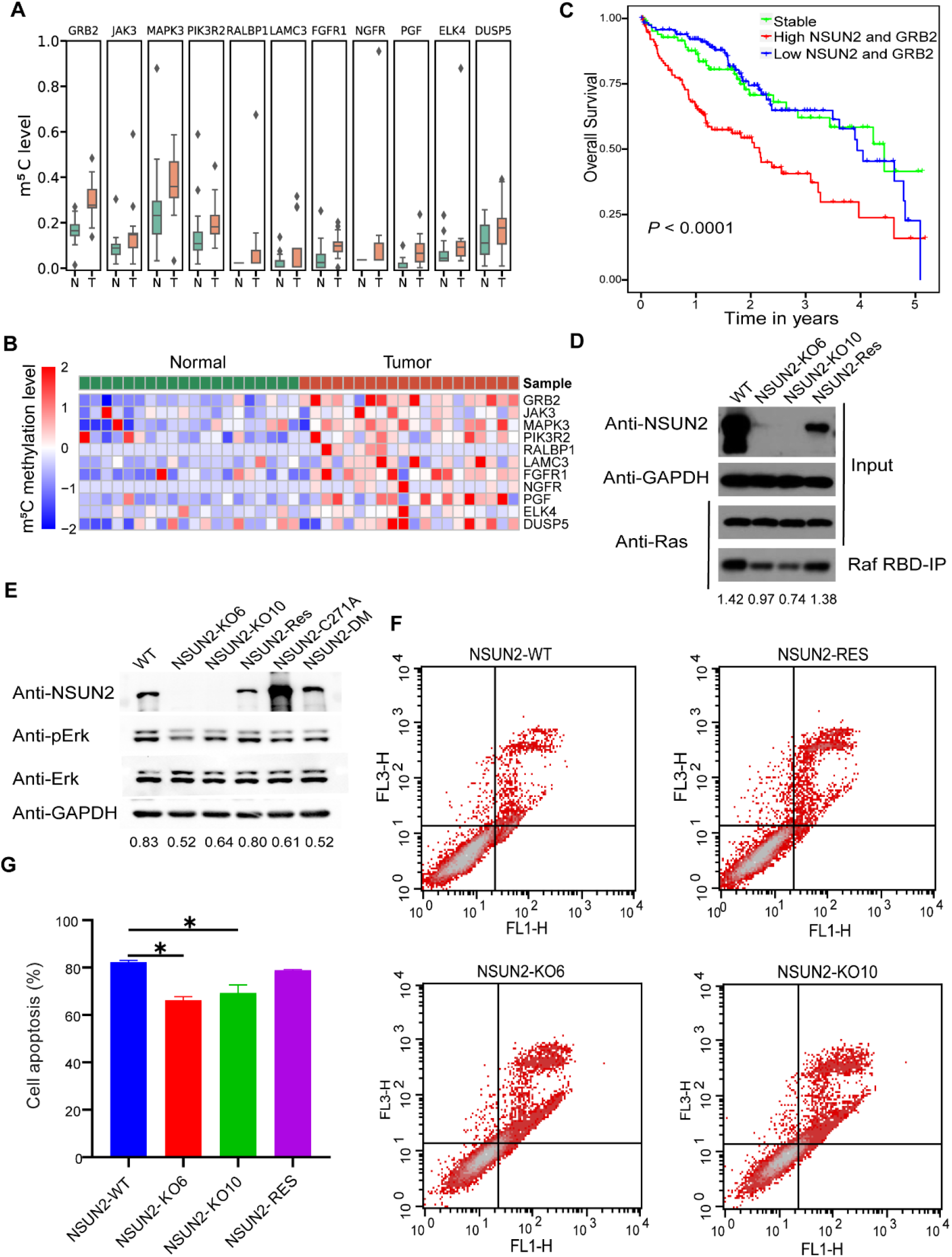
NSUN2 affects the sensitivity of HCC cells to sorafenib by regulating the activity of the Ras pathway. **A**. The mRNA m^5^C modification of the Ras pathway-related genes was analyzed at an integral level. **B**. A heatmap shows differential m^5^C modification of these mRNAs in HCC tissues and the adjacent tissues. **C**. The overall survival analysis indicated that high expression of NSUN2 and GRB2 was correlated with the worst prognosis in HCC patients (*P* < 0.0001). **D**. Ras activity was detected in wild-type QGY-7703 cells (WT), NSUN2-deficient cells (NSUN2-KO6/NSUN2-KO-10), NSUN2-reconstituted cells (NSUN2-Res). **E**. Western blotting of Erk and phosphorylated-Erk (p-Erk) in NSUN2 knockout cells and reconstituted cells, GAPDH was used as a reference control. **F**. Sorafenib treatment and flow cytometry analyses the apoptosis of QGY-7703 cells, when NSUN2 was knockout or rescued. **G**. The statistical analysis of the apoptosis ratio of Figure 4F, \*\**P* < 0.01. Statistical significance was calculated by student’s t-test, mean ± SE.

Initially, we constructed two NSUN2-deficient (KO-6/KO-10) cell lines and one NSUN2-reconstituted stable cell line (QGY-7703). The identification of NSUN2 knockout at the genome level and the mRNA expression level are shown in Figure S3. By western blot analysis, we found that the level of active Ras protein in NSUN2 knockout cells was significantly decreased, which could be rescued by reconstitution of NSUN2 (Figure 4D). The level of phosphorylated-Erk (p-Erk) was an important indicator of the activity of the Ras pathway. We found that p-Erk decreased in HCC cells without changing the Erk protein level in NSUN2 knock-out cells, and rescued by reconstitution of wild NSUN2. The changing of p-Erk level can not be rescued by reconstitution of mutant NSUN2 (Figure 4E).

Sorafenib is a phosphokinase inhibitor and is widely used in the systemic therapy of HCC. Sorafenib inhibits the phosphorylation of Raf, which is a downstream target of active Ras. We had previously shown that NSUN2 could regulate the activity of Ras protein. We investigated whether NSUN2 affects the sensitivity of sorafenib in HCC cell lines. For this, we performed flow cytometry analysis of the apoptotic HCC cells in the NSUN2 knockout group and the control group under sorafenib stress. As expected, the proportion of apoptotic HCC cells treated with sorafenib was significantly higher, compared to the proportion of apoptotic cells in the control group (Figure 4F). Data statistics are shown in Figure 4G. Similar results were obtained for the other HCC cells (Figure S4A & S4B). In addition, we observed that knockdown of NSUN2 did not increase the apoptotic rate of QGY-7703 cells but affected that of Hun7 cells. The increased sensitivity of these cell lines to sorafenib is consistent, when NSUN2 is knocked down. (Figure S4C). Additionally, we found that the downregulation of NSUN2 was associated with cell cycle arrest (Figure S4D & S4E). In summary, NSUN2 affects the sensitivity of HCC cells to sorafenib by regulating the activity of the Ras pathway.

## Discussion

In recent years, many RNA modifications have been identified [27]. As an essential epigenetic modification of RNA, m^5^C participates in different regulatory mechanisms and biological functions, especially in cancers. In this study, the distribution characteristics of the m^5^C modification in HCC were studied. We discovered high levels of m^5^C modification and NSUN2 expression in HCC. The hypermethylated target genes (*GRB2, AATF, RNF115*, and so on) can participate in the carcinogenic pathway. NSUN2 affects the sensitivity of HCC cells to sorafenib by regulating the activity of the Ras pathway.

NSUN2 is highly expressed in multiple tumor types, such as gastric cancer and esophageal squamous cell carcinoma [18, 28, 29]. Here, we found that NSUN2 was overexpressed in HCC. Moreover, NSUN2 in HCC tissues was strongly correlated with the high methylation and expression of target genes, including *GRB2, RNF115, AATF, ADAM15, RTN3*, and *HDGF*. Li et al. showed that NSUN2 coordinates with lin-28B, a novel m^5^C recognition protein, to catalyze the m^5^C modification of GRB2 and stabilize its mRNA expression. High levels of GRB2 promote the activation of the PI3K/AKT and Ras pathways in esophageal squamous cell carcinoma [30]. We demonstrated that NSUN2 inhibited the Ras activation and decreased the p-ErK level in HCC, which led to the increased sensitivity of HCC cells to sorafenib.

Studies have shown that nascent RNA with m^5^C modification can regulate chromatin structures and recruit transcription factors. The m^5^C modification-mediated complex leads to 5-azacitidine resistance in leukemia cells, which provides new insight into the treatment of leukemia [31]. In our study, because of the critical role of NSUN2 in regulating the m^5^C modification and the expression of mRNAs related to the Ras pathway, the sensitivity of HCC cells to sorafenib was increased, which has great significance for the treatment of HCC patients. RNA epigenetics, especially m^5^C, can potentially regulate drug sensitivity.

Overall, we found the m^5^C modification in HCC at a single nucleotide resolution landscape and verified the correlation between m^5^C hypermethylated genes and HCC tumor characteristics. NSUN2 was found to be involved in various tumor-related cell processes, including affecting proliferation, apoptosis, and sorafenib sensitivity in HCC cells. Our study provided novel mechanisms for the effect of RNA epigenetic modification on HCC progression, which might help to discover more effective HCC treatment targets and strategies.

## Materials and methods

### Cell lines and tissues

Human HCC cell lines (QGY-7703, Huh 7) were cultured in Dulbecco’s Modified Eagle Medium (DMEM) supplemented with 10% fetal bovine serum and 1% penicillin/streptomycin in 5% CO_2_ at 37 °C. For sensitivity analysis of sorafenib, QGY cells were seeded in six-well plates and treated with 10 µM sorafenib for 24 h. The Huh7 cells were treated with 8 µM sorafenib for 24 h.

### Plasmids, antibodies, and real-time PCR primers

PLKO.1-shNC, pLKO.1-shNSUN2, psPAX2, and pMD2 were used for NSUN2 knockdown in HCC cells. The sequence of shNSUN2 was 5-CCGGGCTGGCA CAGGAGGGAATATACTCGAGTATATTCCCTCCTGTGCCAGCTTTTTG-3. The sequence of siNSUN2 was: 5-CACGUGUUCACUAAACCCUAUTT-3.

The antibodies used in this study were antibody-NSUN2 (CST, 44056S), antibody-GAPDH (Servicebio, GB12002), antibody-pErk (CST, 4370s), and antib ody-Erk (CST, 4695s). Real-time PCR primers for target gene quantification we re used in this study, including RNF115 (forward 5-CGGCAGTCGGATAGACA ATAC-3, reverse 5-TGTCAGGACGAGAACTTCCTC-3), GRB2 (forward 5-CTG GGTGGTGAAGTTCAATTCT-3, reverse 5-GTTCTATGTCCCGCAGGAATATC-3), ADAM15 (forward 5-CCCTGAATGTACGAGTGGCAC-3, reverse 5-GGAGGA AGTTTTCGAGGGTGA-3), and AATF (forward 5-TCAGCCTCCCTCTTGGAC A-3, reverse 5-TCATCAGACGATCCTGGCAGA-3). NSUN2 (forward 5-GGTAT CCTGAAGAACTTGCC-3, reverse 5-ATCTTATGATGAGGCCGCA-3). GAPDH (forward 5-CGCTCTCTGCTCCTCCTGTTC-3, reverse 5-ATCCGTTGACTCCGACCTTCAC-3).

### RNA-BisSeq and RNA-seq for HCC tissues and adjacent tissues

Tissues were frozen in liquid nitrogen and broken using Qiagen Tissue Lyser II. Then, 1 mL TRIzol was added to the broken tissues, and total RNA was extracted with chloroform-isopropyl alcohol. Next, the HCC mRNAs were enriched using the Dynabeads mRNA purification kit (Ambion, 61006) protocol, and samples were treated with DNase (Thermo, AM22222) at 37 °C for 20 min to remove genomic DNA. After DNase treatment, RNA was fragmented by a fragmentation reagent (Ambion, AM8740), and then the alcohol method was used to precipitate the samples.

After alcohol precipitation, 10 ng of mRNA samples were taken for the transcription library, and 100–200 ng of mRNA samples were taken for bisulfite treatment, according to an earlier published method [32]. Finally, we used the KAPA Stranded RNA-Seq Library Preparation Kit (KAPA, KR1139) for library construction. Sequencing was performed on an Illumina HiSeq PE150 sequencing system with paired-end 150 bp read length.

### UHPLC–MS/MS analysis

The UHPLC–MS/MS analysis was performed by a previously reported method [13]. Total RNA or mRNA (100–200 ng) was extracted from the QGY-7703 and SMMC-7721 cells, which were digested with 0.1 U Nuclease P1 (NEB, M0660) and 1.0 U calf intestinal alkaline phosphatase (Invitrogen, 18009019) at 37 °C overnight. Then, the mixture was filtered through a 3 K Omega Membrane tube (PALL, OD010C35). Finally, we detected rm^5^C, rC, rU, rG, and rA using UHPLC–MS/MS.

### Immunohistochemistry (IHC)

The tissues were fixed with 5 mL of formaldehyde fixative solution. Then, they were dehydrated by adding molten paraffin wax at 58 °C. Tissues were cut into 15 µm sections using a rotary microtome, suspended in a water bath at 56 °C, and mounted onto gelatin-coated histological slides. The slides were dried overnight at room temperature. Then, we performed immunohistochemistry analysis. The samples were incubated with NSUN2 antibody (1:100) overnight at 4 °C. Finally, the expression of NSUN2 in HCC tissues was visualized under a microscope using bright-field illumination.

### Ras activation assay

RAS activity was analyzed with a Ras Activation Assay Biochem Kit (Cytoskeleton, BK008). The QGY cell lines containing the control group, the NSUN2-KO6 group, the NSUN2-KO10 group, and the NSUN2-Res group were prepared in advance, and equal concentrations of cells were collected and spread in a six-well plate. After 24 h, 500 µL cell lysate was added to each well and centrifuged at 4 °C and 10,000 rpm for 2 min, and the supernatant protein was collected. The Bradford protein quantification kit was used to quantify the protein, and each group was diluted with cell lysate to equal volume and density. Then, 20 µL of whole-cell lysate was added to 5 µL of 5 × SDS loading buffer, and the sample was boiled at 95 °C for 10 min as an input sample. The remaining samples were added with the same amount of RAF-RBD beads and rotated at 4 °C for 1 h. The beads were collected at 4 °C and centrifuged at 5,000 g for 1 min. Then, 90% of the supernatant was removed, and the beads were cleaned three times with 500 µL of Wash Buffer. Finally, 1 × SDS loading buffer was added, and the sample was boiled at 95 °C for 10 min as an IP sample. The samples were subjected to Western blot electrophoresis, and the anti-Pan RAS antibody was used to quantitatively identify the active RAS through co-immunoprecipitation using RAF-RBD beads in each group of cells.

### Flow cytometry analysis

The NC (shControl) group and the shNSUN2 group cells were seeded in a six-well plate. The cell confluence reached 80% through overnight culture. Then, the cells were treated with sorafenib for 24 h. The cells were harvested and washed once with precooled PBS. According to the protocol of the Dead Cell Apoptosis Kit (Thermo, V13241), 5x annexin-binding buffer was diluted to 1× with deionized water, and the PI staining solution was diluted to 100 µg/mL. After the buffer was prepared, the cells were resuspended in 100 µL 1× annexin-binding buffer, 5 µL Alexa Fluor 488-annexin V, and 1 µL PI (100 µg/mL). The cells were incubated for 15 min at room temperature. Then, 400 µL of 1× annexin-binding buffer was added and gently mixed before flow cytometry analysis. All the experiments were repeated at least three times.

### RNA-seq analysis

The raw data were trimmed for adaptors by the Cutadapt software, and low-quality bases were removed by the Trimmomatic software [33, 34]. The filtered clean reads were mapped to the hg19 genome with hisat2 [35]. The HTSeq software was used to count reads mapped to each Ensembl gene [36]. Differentially expressed genes were calculated using DESeq2 [37]. The differential fold change cutoff was 1.2, and the FDR value cutoff was 0.05.

### RNA-BisSeq bioinformatics analysis

The Cutadapt and Trimmomatic software were used to trim adaptors and remove low-quality bases [33, 34]. The clean reads were mapped to the hg19 genome by meRanGh from meRanTK[38].

The m^5^C sites were called by meRanCall from meRanTK. The luciferase spike-in conversion rates were evaluated to be over 99%. The sample-credible m^5^C sites satisfied coverage depth ≥ 30, methylated cytosine depth ≥ 5, and methylation level ≥ 0.1. The differential m^5^C methylation analysis criteria comprised coverage ≥ 10 for all samples and were used to compare methylation levels between tumor and normal samples. The differential m^5^C sites were defined as follows: mean m^5^C level difference ≥ 0.05 (tumor and normal samples) and p-value < 0.05 (Wilcoxon test). The m^5^C sites were annotated using bedtools intersectBed [39].

### Pathway analysis

Hypermethylated and hypomethylated genes were used as input for DAVID (https://david.ncifcrf.gov/).

### Statistical analysis

Data were analyzed using the Python and GraphPad Prism 8 software. Two-way analysis of variance and Student’s t-test were performed to determine statistical significance. The error bars, when present, represent the mean ± SD. The experiments were repeated at least three times independently. Statistical significance was considered at *P* < 0.05.

## Supporting information

Supplementary table 1. The list of mRNA m5C sites in HCC.

## Ethical statement

The 20 paired samples of hepatocellular carcinoma tissues and adjacent tissues in this study were collected from the biobank of the First Affiliated Hospital of Zhengzhou University. Written informed consent was obtained from all patients, and an ethical review was passed. Approval No. 2022-KY-0005-002.

## Data availability

The raw sequence data reported in this paper have been deposited in the Genome Sequence Archive [40] (Genomics, Proteomics & Bioinformatics 2021) in the National Genomics Data Center, Beijing Institute of Genomics, Chinese Academy of Sciences / China National Center for Bioinformation (GSA: HRA001101), which is publicly accessible at https://ngdc.cncb.ac.cn/gsa.

## CRediT authors statement

**Dan Song:** Methodology, Validation, Investigation, Writing - Original Draft, Writing - Review & Editing. **Ke An:** Formal Analysis, Data Curation, Writing – Original Draft Preparation.Writing - Review & Editing. **Wen-Long Zhai:** Resources, Methodology, Investigation, Writing. **Lu-Yao Feng:** Validation, Investigation. **Ying-Jie Xu:** Validation, Investigation. **Ran Sun:** Validation. **Yue-Qin Wang:** Methodology, Investigation. **Yun-Gui Yang:** Methodology, Supervision, Project Administration. **Quan-Cheng Kan:** Conceptualization, Project administration, Funding Acquisition. **Xin Tian:** Conceptualization, Supervision, Funding Acquisition, Project Administration, Writing - review & editing. All the authors have read and approved the final manuscript.

## Acknowledgments

We would like to thank the patients who generously donated HCC tissues that made this research possible. We also thank Professor Jiaxue Wu from Fudan University for providing HCC cell lines. This study was financially supported by grants from the National Natural Science Foundation of China (Grant Nos. 32170594 and 31870809), the Province and Ministry Coconstruction Major Program of Medical Science and Technique Foundation of Henan Province (No. SBGJ202001007), and the Special Fund for Young and Middle School Leaders of Henan Health Commission (No. HNSWJW-2020017).

## Figure legends

**Supplementary Figure 1.**
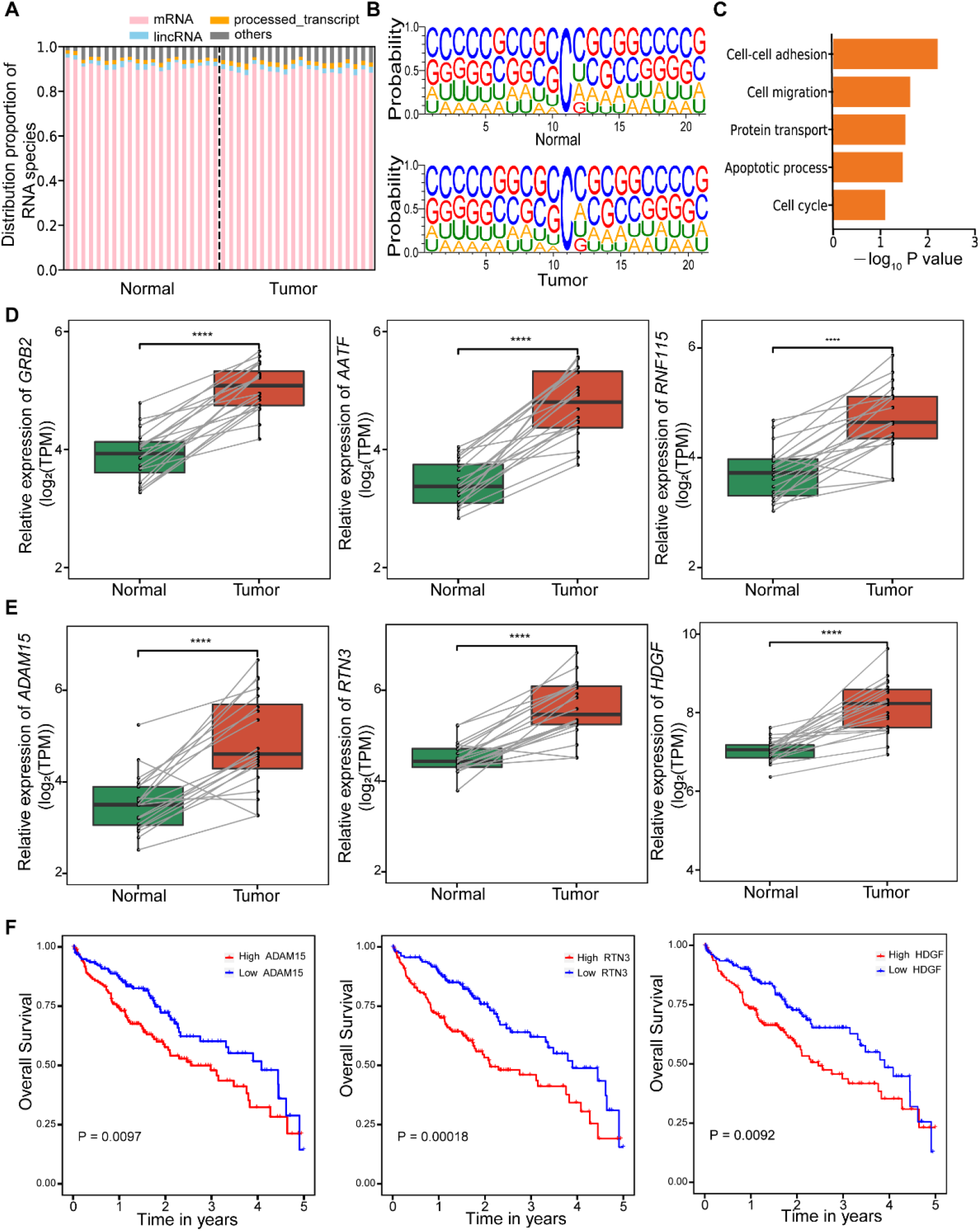
Distribution characteristics of m^5^C in HCC and expression pattern of target genes. **A**. The type and proportion of RNA in the sequencing library. **B**. The sequences are proximal to the mRNA m^5^C sites in HCC tissues and the adjacent tissues. **C**. The m^5^C hypermethylated and highly expressed genes in HCC tissues were involved in different cellular processes, determined by the GO functional analysis. **D** & **E**. The expression of target genes, including *GRB2, AATF, RNF115, ADAM15, RTN3*, and *HDGF*, was higher in HCC tissues than that in the adjacent tissues, determined by transcriptome analysis, \*\*\*\**P*< 0.0001. **F**. The overall survival analysis showed that the higher expression of ADAM15, RTN3, and HDGF was correlated with a poor prognosis in HCC patients; ADAM15 (\**P* = 0.0097), RTN3 (\*\**P* = 0.00018), HDGF (\**P* = 0.0092).

**Supplementary Figure 2.**
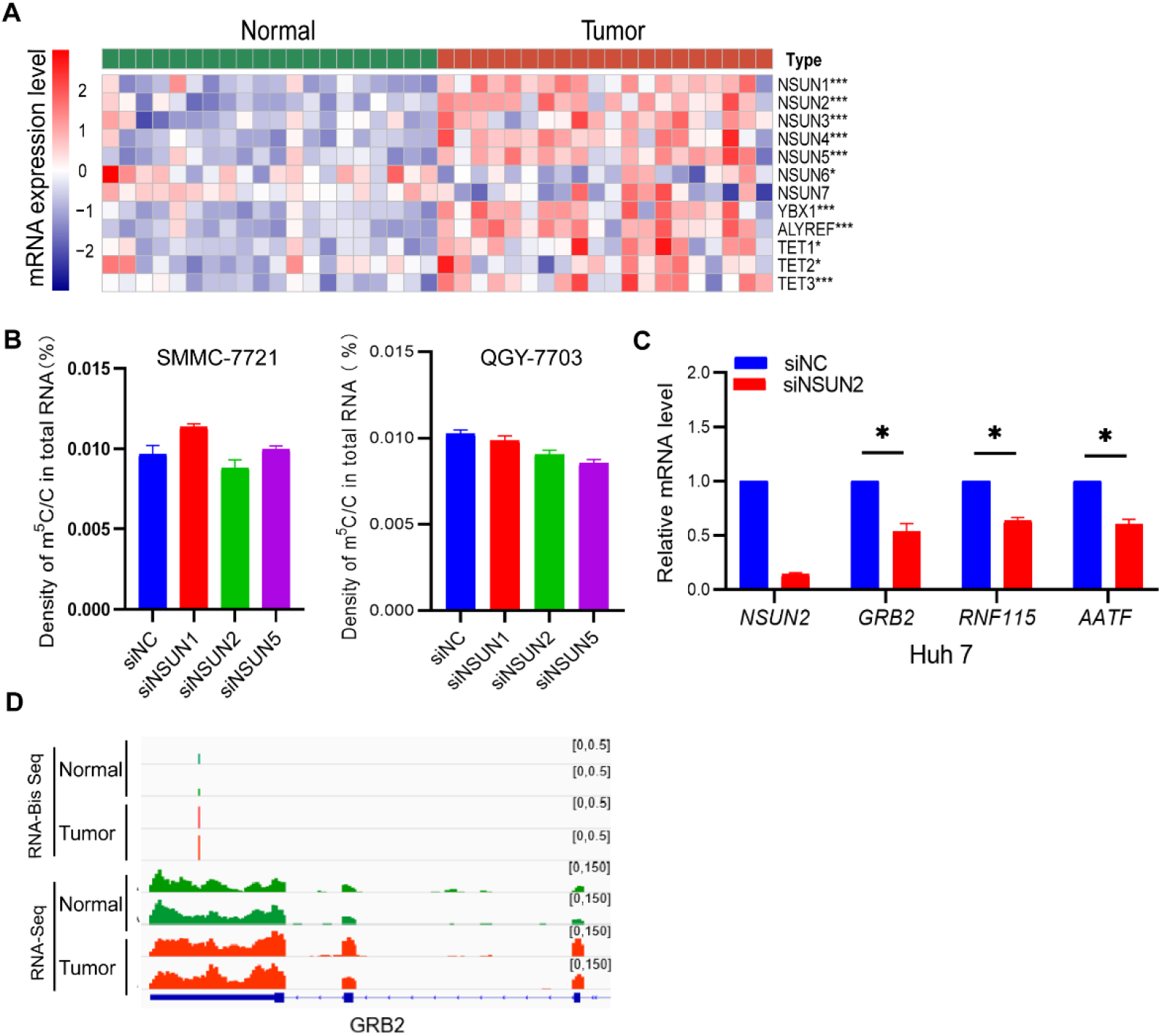
NSUN2 regulates downstream target genes expression, especially GRB2. The mRNA expression of m^5^C writers and readers in HCC tissues was higher than that in the adjacent tissues, determined by the transcriptomic analysis, ***, *P* < 0.001. In HCC cell lines, the UHPLC–MS/MS analysis showed that downregulation of NSUN2 did not change the density of m^5^C/C in total RNA. **C**. Real-time PCR analysis showed that the mRNA expression levels of *GRB2, RNF115*, and *AATF* were significantly lower in Huh7 NSUN2 knockdown cells, \**P* < 0.05; \*\**P* < 0.01. **D**. The m^5^C modification in the 3’UTR and the expression of GRB2 mRNA were analyzed by IGV visualization in HCC tissues and the adjacent tissues.

**Supplementary Figure 3.**
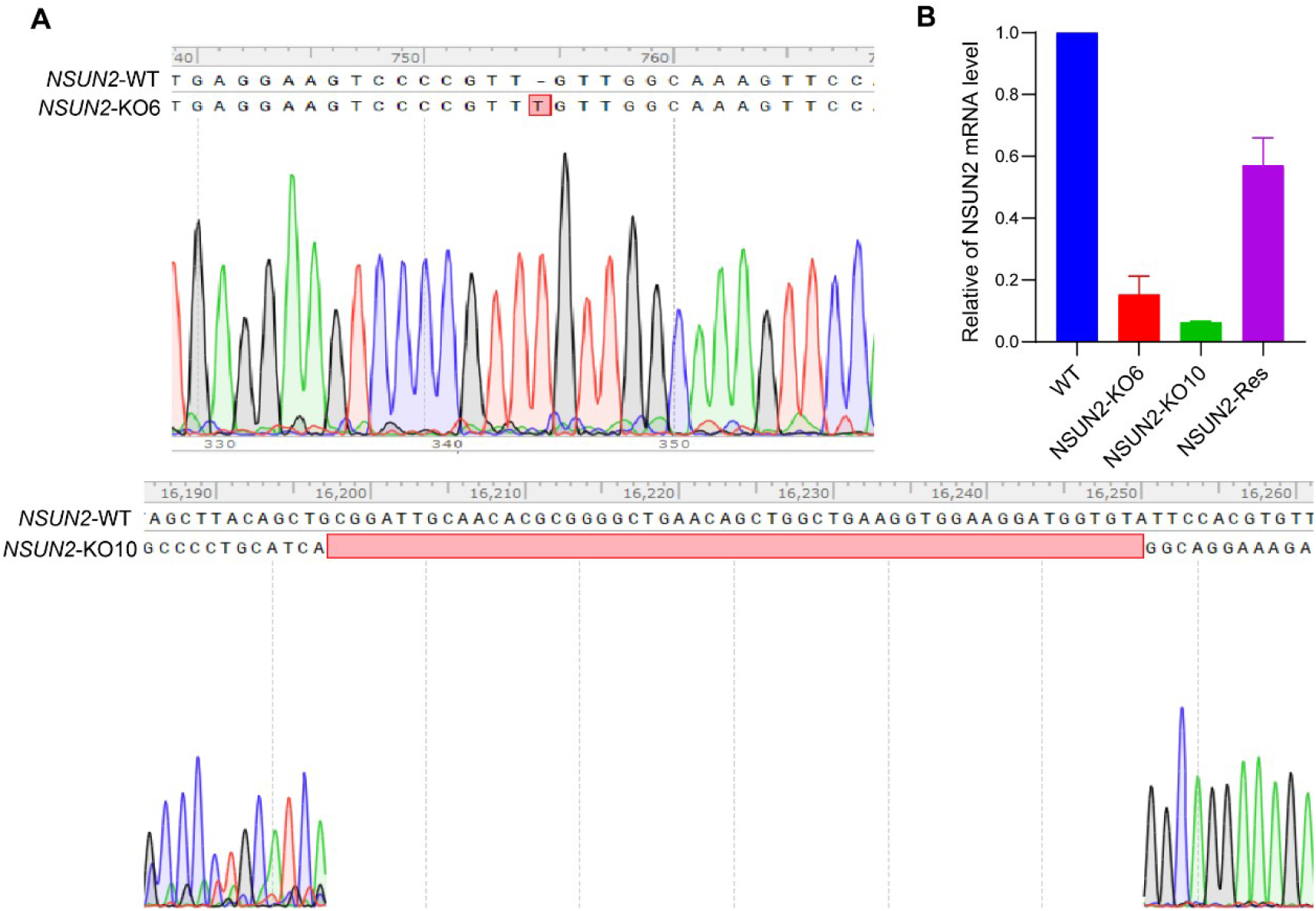
Identification of NSUN2 knockout efficiency at genome level and transcriptome level. **A**. NSUN2 knockout characteristics were identified by Sanger sequencing in the QGY-7703 cell genome. **B**. The mRNA expression levels of NSUN2 in NSUN2 knockout cells (KO-6/KO-10) and NSUN2 reconstituted cells were evaluated by real-time PCR.

**Supplementary Figure 4.**
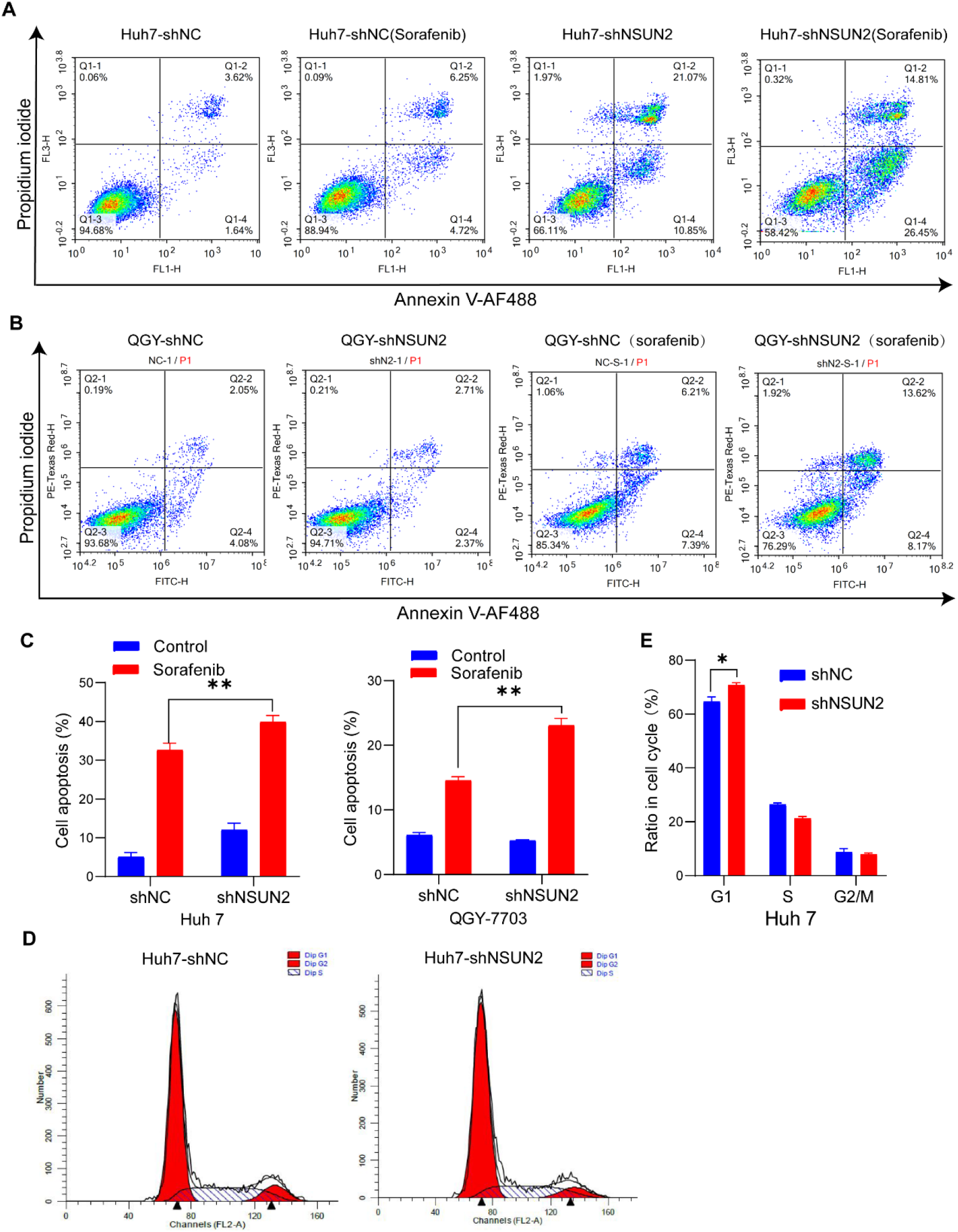
Apoptosis and cycle analysis of NSUN2 on HCC cells. **A & B**. After sorafenib treatment, flow cytometry analyses of the apoptosis of HCC cells rate when NSUN2 knockdown. The statistical analyses of the apoptosis ratio are shown in **C**, \*\**P* < 0.01. **D**. Flow cytometry analyses of the cell cycle of Huh 7 cells. These cells were arrested in the G1 phase when NSUN2 was knocked down. The statistical analyses of the arrest ratio are shown in **E**, \**P* < 0.05. Statistical significance was calculated by student’s t-test, mean ± SD.

**Supplementary table 1. The list of mRNA m**^**5**^**C sites in HCC**.

